# Discovery of Small Molecule CHI3L1 Inhibitors by SPR-Based High-Throughput Screening

**DOI:** 10.1101/2025.08.24.671973

**Authors:** Longfei Zhang, Hossam Hammouda Nada Hammouda, Moustafa T. Gabr

## Abstract

Chitinase-3-like 1 (CHI3L1) is a secreted glycoprotein implicated in carcinogenesis and tumor immune evasion. Elevated CHI3L1 expression is frequently detected in cancer patients, highlighting it as a promising therapeutic target. To overcome the limited availability of small molecule CHI3L1 inhibitors, we established a surface plasmon resonance (SPR)–based high-throughput screening platform and applied it to a focused chemical library of small molecules. Primary screening identified seven hits, with compounds **1-4** and **1-7** validated as CHI3L1 binders (*Kd* = 10.4 ± 1.0 μM and 7.40 ± 0.78 μM, respectively). Both compounds disrupted the CHI3L1–galectin-3 interaction in AlphaLISA assays and engaged the CHI3L1 binding pocket in docking and molecular dynamics (MD) simulations. Importantly, functional evaluation in a multicellular 3D glioblastoma (GBM) spheroid model demonstrated that compound **1-7** potently reduced spheroid viability and inhibited STAT3 phosphorylation, outperforming both compound **1-4** and the known CHI3L1–STAT3 disruptor hygromycin B (HB). These findings validate SPR as a robust primary screening platform for CHI3L1 and demonstrate that the identified small molecule binders exert functional activity in a physiologically relevant multicellular GBM spheroid model.

## INTRODUCTION

Even though lacking of enzymatic activity characteristic of other members in the glycoside hydrolase 18 family,^1–3^ as a result of the mutation in its catalytic domain, chitinase-3-like 1 (CHI3L1) can still stimulate multiple signaling pathway (ERK1/2, AKT, WNT/β-catenin, NF-κB, and STAT3) via interacting with different receptor proteins, such as IL-13Rα2, galectin-3, and TMEM219,^4^ thus involved in various biological process, including inflammation, pathogen defense, and wound healing.^2, 5–7^ However, in recent decades, accumulating evidence revealed that CHI3L1 also plays vital roles in carcinogenesis, and elevated CHI3L1 expression is frequently detected in both serum and tumor tissue of patients with various cancers, including non-small cell lung cancer, hepatocellular carcinoma, and glioblastoma (GBM).^8–11^ Importantly, clinical studies have demonstrated a positive correlation between serum CHI3L1 levels and cancer stage, with the high expression generally associated with poor prognosis and reduced survival.^12–15^ Consequently, CHI3L1 has attracted increasing attention as both a diagnostic biomarker and potential therapeutic target in cancer treatment.

During carcinogenesis, CHI3L1 recruits immune cells, such as neutrophils and macrophages, into the tumor region and stimulates them to release pro-inflammatory cytokines (IL-6, IL-8, IL-1β, and TGF-β) to sustain a chronic inflammatory environment, which promotes cancer cell proliferation and migration while inhibiting apoptosis.^16, 17^ CHI3L1 also drives the recruitment of tumor-associated macrophages (TAMs) and polarizes them into an immunosuppressive M2-like phenotype via an IL-13Rα2-relevant signaling pathway, creating an immunosuppressive microenvironment and facilitating tumor immune evasion.^14, 18^ In particular, STAT3 has been implicated as a critical mediator of this immune suppression, linking CHI3L1 activity to macrophage polarization and the inhibition of antitumor immunity.^14, 18^ In 2021, after intravenously injecting anti-CHI3L1 antibody at a dosage of 0.5 mg/kg body-weight on multiple tumor-bearing mice models (LLC cell, A549 cell, and B16-F10 cell), Yu et al. observed a dramatic decrease in tumor volume, tumor weight or metastasis number, accompanied with a reduction in serum and tumor CHI3L1 expression level compared to the control group.^14^ Following quantitative assay using RT-qPCR and western blot revealed a significant decrease in the expression of M2 markers (ARG1 and CD206) in both tumor and metastasis after anti-CHI3L1 treatment compared to the control group, while iNOS and CD86 protein (M1 markers) remain unaffected, indicating the shift towards M1-like phenotype.^14^ Moreover, CHI3L1 was also reported to impair the antitumor immunity by skewing the Th1/Th2 balance towards Th2 through IFN-γ-related signaling.^16, 19^ Besides indirectly sustaining tumor growth via modulating the microenvironment, CHI3L1 could be secreted directly by cancer cells to promote the proliferation and migration. In 2012, Kawada et al. transfected SW480 cells with a CHI3L1 vector and CHI3L1-specific miRNA to build up a CHI3L1 overexpression and silenced phenotype, respectively. In the following assays, compared to the control group, an obvious increase in cell proliferation and migration was observed in CHI3L1 overexpression cells, while silenced phenotype SW480 cells showed a reduction in both properties.^20^ In the following *in vivo* experiment, after 4 weeks post-subcutaneous inoculation, CHI3L1 vector-transfected HCT116 cells induced tumors with 3.5-fold in volume, accompanied by elevated macrophage infiltration compared with the control cells, highlighting the viral role of CHI3L1 in tumorigenesis.^20^

Although CHI3L1-inhibition has demonstrated therapeutic promise on animal models, as the development remains at an early stage, inhibition reagents used in *in vivo* experiments are still limited to antibodies, while scarcely molecular inhibitors have been reported. Compared to antibodies, small-molecule drugs demonstrated distinct advantages in tissue penetration, oral bioavailability, manufacturing cost, and transportation. Moreover, benefiting from the simple chemical structure, the problematic immune-related adverse events commonly associated with antibody treatment could be avoided in small molecule medicines, and the easy chemical structure modification facilitates the following pharmacokinetics optimization. In 2024, Czestkowski et al. reported two molecular inhibitors, compounds **30** and **36**, which were identified using an AlphaScreen-based high-throughput screening (HTS) targeting the interaction between CHI3L1 and a fractionated heparan sulfate polymer (HSIII).^21^ Both compounds demonstrated potent inhibition abilities with IC_50_ values of 37 nM and 26 nM, respectively.^21^ **K284-6111** was identified by Prof. Hong’s laboratory through a structure-based virtual screening, which is one of the most developed molecular CHI3L1 inhibitors, and demonstrated promising therapeutic performances on both cancer and Alzheimer’s disease animal models.^22–24^ On both B16F10 and A549 metastasis mice model, after intravenously administration of **K284-6111** at a dosage of 0.5 mg/kg body-weight at 3-day intervals for 3 or 8 weeks, respectively, a significantly reduction in metastatic nodule numbers were observed compared to the vehicle administration group, accompanied with a decreased CHI3L1 expression level in tumor region, highlighting the therapeutic potential of **K284-6111** in cancer treatment.^22^

Although several CHI3L1 molecular inhibitors have been reported, most displayed relatively weak binding affinities or lacked robust quantitative affinity data, limiting subsequent structure optimization and clinical transformation. To address this issue, developing a novel HTS platform capable of identifying ligands with stronger affinities is both urgent and necessary. Surface plasmon resonance (SPR) is an optical technique designed to measure the interaction between the target protein and its ligand in real time by detecting refractive index changes at the surface of a sensor chip. Compared with other platforms commonly used in HTS, including TRIC, FRET, and AlphaLISA, SPR offers distinct advantages in precision, sensitivity, and reliability. Since the direct signal readout is the shift of incident polarized light, it is less susceptible to interference from fluorescence background, which makes the screening of fluorescent molecules possible, while it has always been a problematic issue in most fluorescence-based HTS platforms. More importantly, SPR provides detailed kinetic information even at a single concentration, enabling the early identification and exclusion of non-specific or weak binders, thereby reducing experimental workload and reagent costs in downstream assays.

In this study, to evaluate the feasibility of SPR as a CHI3L1 HTS platform and discover novel molecular binders, 704 molecules from the Discovery Diversity Set (DDS-10, Enamine, Kyiv, Ukraine) were screened on the Biacore™ 8K SPR system (Cytiva, Marlborough, MA, USA). After the single-concentration binding assay, 2 hits were highlighted with high occupancy and favorable binding kinetic, corresponding to a hit rate of 0.28% (**Figure 1a**). Subsequent binding affinity measurement validated both compounds **1-4** and **1-7** as potent CHI3L1 binders, and the binding affinities were calculated as 10.4 ± 1.0 μM and 7.40 ± 0.8 μM, respectively. Moreover, in the AlphaLISA-based inhibition assay, both compounds showed inhibitory activities against the interaction between CHI3L1 and galectin-3, and the IC_50_ values were measured as 23.5 ± 2.0 μM and 15.4 ± 4.7 μM, respectively.

**Figure 1.**
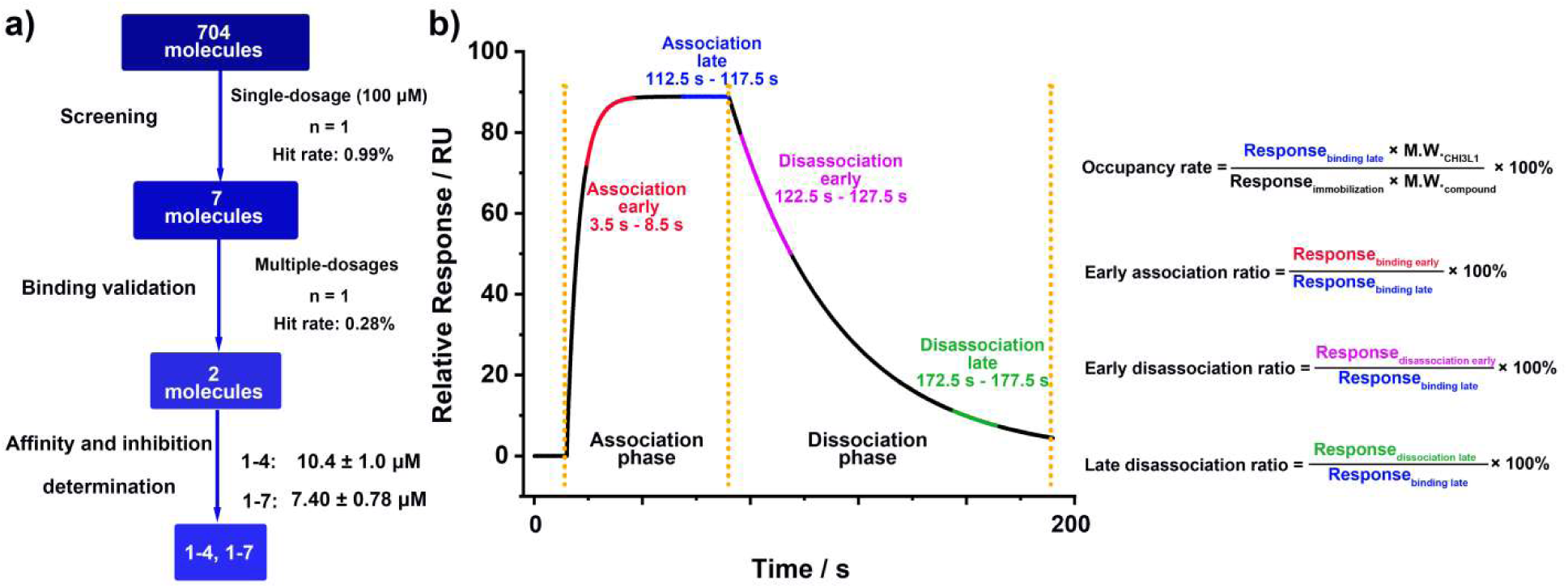
SPR-based HTS workflow and sensorgram analysis. **a)** Schematic illustration of the overall SPR-HTS pipeline used to identify CHI3L1 binders. A focused library of 704 molecules was screened at a single concentration (100 μM), yielding seven preliminary hits. Subsequent multi-concentration validation reduced this to two confirmed binders, compounds **1-4** and **1-7**. **b)** Representative sensorgram illustrating association and dissociation phases, with time windows for early and late association/dissociation highlighted in red, blue, purple, and green, respectively. Quantitative parameters used for hit evaluation are shown on the right: occupancy rate, early association ratio, early dissociation ratio, and late dissociation ratio.

## RESULTS AND DISCUSSION

### SPR-based Primary Screening

For discovering novel CHI3L1 molecular binders, 704 molecules from the DDS library were screened using SPR. In the primary screening, to save workload and reagents, the binding check was performed at a single concentration (100 μM). CHI3L1-His protein was freshly immobilized on both eight channels of a Series S Sensor Chip CM5 (29104988, Cytiva, Marlborough, MA, USA) by covalent coupling, achieving immobilization levels of 3000 - 3600 Response Unit (RU). 100 μM compounds in the running buffer (supplemented with 2.5% DMSO) were injected over the sensor chip at a flow rate of 30 μL/min for 120 s, followed by a 600 s dissociation phase.

During screening, the compound injections were performed in sequence, and a 30 s regeneration phase (regeneration buffer: 10 mM sodium acetate, 250 mM NaCl, pH 4.0) was operated between two injections to wash off any residual from the previous candidate. Running buffer with 2.5% DMSO served as the negative control and was injected every 24 cycles, and the solvent correction was carried out every 48 cycles. A total 96 compounds were tested per channel, and no protein re-immobilization was required throughout the screening.

In SPR, occupancy rate reflected the binding extent under the 1:1 binding assumption at the tested concentration and is commonly used as a rough indicator of binding strength (**Figure 1b**). However, this parameter cannot reliably distinguish non-specific interactions. In contrast, the sensorgram, which plots response over time, provides detailed kinetic information and is a powerful tool for assessing binding specificity. Nevertheless, manually inspecting sensorgram shapes for each compound is labor-intensive and subjective, particularly in HTS campaigns involving hundreds of candidates. To enable a more quantitative and efficient evaluation, we defined three sensorgram-based parameters: (i) early association ratio = (Response_association early_ / Response_association late_) × 100 %, reflecting the speed of association; (ii) early dissociation ratio = (Response_dissociation early_ / Response_association late_) × 100 %, indicating the initial dissociation rate, and (iii) late dissociation ratio = (Response_dissociation late_ / Response_association late_) × 100 %, representing the overall dissociation level (**Figure 1b**).

In the primary screening, 33 compounds were excluded due to solubility issues, abnormal response shift, or high reference cell signal (**Figure S1**). Among the remaining 671 compounds, a threshold of > 70% occupancy rate was applied to define primary hits. As shown in **Figure 1a**, 7 compounds meet the criterion, corresponding to a hit rate of 0.99%. To obtain quantitative interaction information, three sensorgram parameters were further calculated. Compounds **1-4** and **1-7** demonstrated a rapid association with early association ratios of 103.3% and 99.5%, respectively, indicating saturation were obtained within 8.5 s post-injection (**Table 1** and **Figure 1b**). Afterwards, the response signals of **1-4** and **1-7** dropped back to ∼ 7.0% of the peak values within 3.5 s of dissociation, suggesting a rapid and complete dissociation for both compounds (**Table 1**). In contrast, compounds **2-4** and **2-5** also reached saturation rapidly (early association ratios of 95.3% and 115.5%, respectively) but dissociated slowly, with early dissociation ratios of 77.5% and 58.7%, respectively (**Table 1**). After 600 s of buffer wash, substantial residual signals remained (late dissociation ratios: 60.2% and 47.0% for **2-4** and **2-5**, respectively), suggesting potential non-specific interactions (**Table 1**). Compounds **4-2**, **5-12**, and **6-1** displayed relatively high occupancy (120.8%, 73.3%, and 71.8%, respectively), but their slow association and dissociation kinetics, along with persistent residual signals (29.8%, 28.4%, and 25.7% of peak responses), indicated non-specific binding behavior (**Table 1**).

**Table 1.**
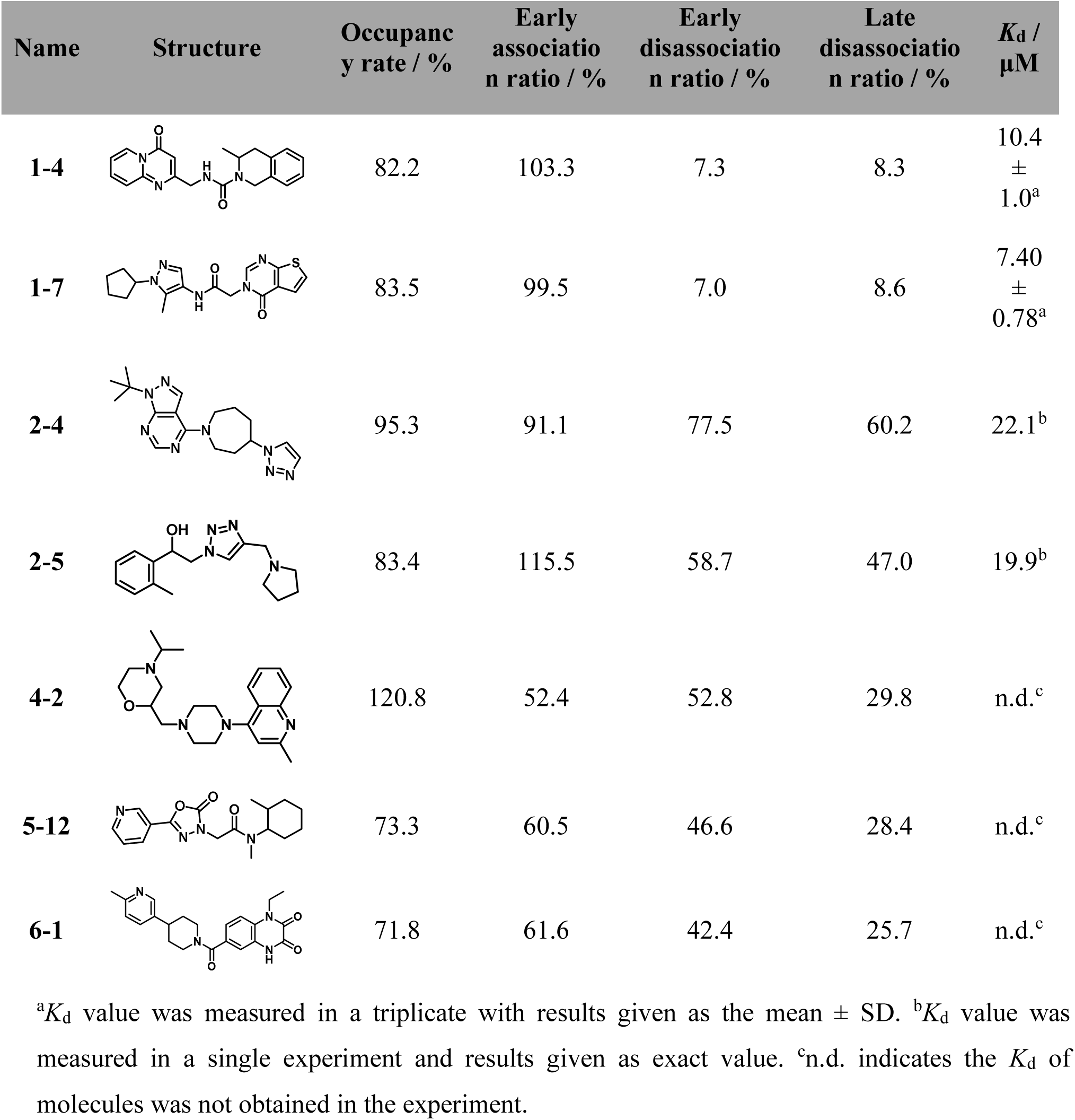
Chemical structures, occupancy rate, early association ratio, early dissociation ratio, late dissociation ratio, and *K*_d_ values of 7 hits identified in primary screening.

Based on the quantitative sensorgram evaluation, **1-4** and **1-7** were inferred as CHI3L1 binders according to the favorable binding kinetics, whereas **2-4** and **2-5** were considered as binders accompanied by potential non-specific binding. Owing to the slow association and dissociation phases, **4-2**, **5-12**, and **6-1** were proposed as non-specific binders. Although such candidates would normally be excluded from further studies, all seven compounds were deliberately retained for subsequent affinity measurements in order to assess the feasibility and reliability of the sensorgram-based screening parameters.

### SPR-based Binding Affinity Measurement

To quantitatively evaluate the binding affinities of previously identified hits towards CHI3L1, a single-cycle kinetic assay (n = 1) was performed on the SPR system. Approximately 3000 RU of CHI3L1 was immobilized on the CM5 sensor chip, and gradient concentrations of the tested were injected at a flow rate of 30 μL/min for 120 s, followed by a 600 s dissociation phase. As shown in **Figure 2**, both **1-4**, **1-7**, **2-4**, and **2-5** exhibited rapid association with CHI3L1, and binding equilibrium was reached at each concentration. Upon dissociation, **1-4** and **1-7** showed a rapid signal decrease to ∼ 10 RU, which then stabilized until the end of the experiment, suggesting the presence of non-specific interaction. In contrast, **2-4** and **2-5** displayed a slow dissociation, with signal continue to decline throughout the dissociation phase. Four compounds displayed moderate to potent binding affinities towards CHI3L1, and the *K*_d_ were calculated as 4.96 μM, 6.39 μM, 23.6 μM, and 10.7 μM for **1-4**, **1-7**, **2-4**, and **2-5**, respectively. Consistent with our early prediction, no clear concentration-dependent response was observed with the increasing concentration for **4-2** and **6-1**, confirming their non-specific interaction with CHI3L1. Owing to the solubility issue, **5-12** could not be reliably analyzed and was excluded from further experiments.

**Figure 2.**
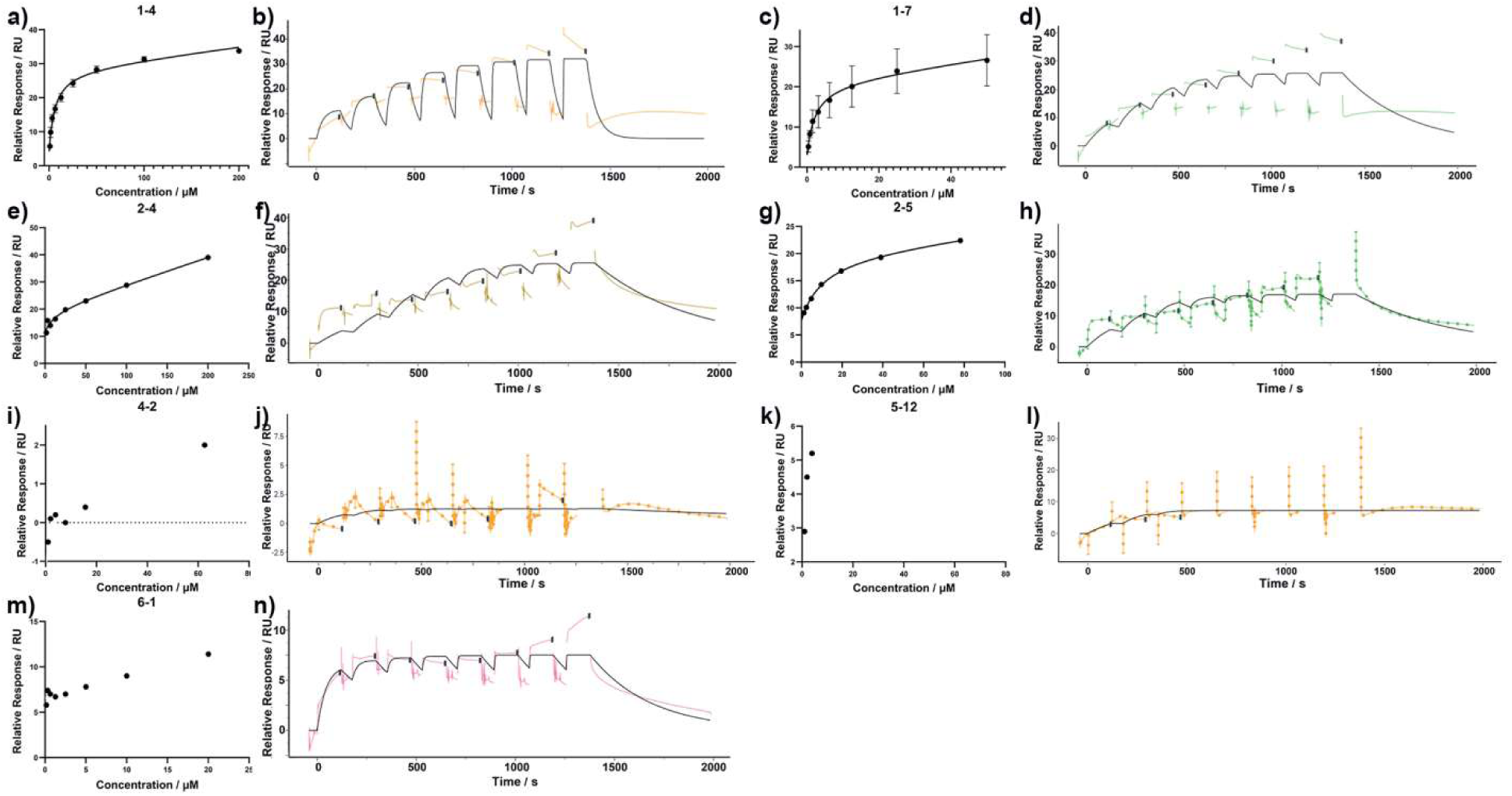
CHI3L1 binding affinity measurement using SPR. SPR-derived plots of relative response versus concentration (left panels) and sensorgrams (right panels) for compounds **1-4** (**a**, **b**), **1-7** (c, d), **2-4** (**e**, **f**), **2-5** (**g**, **h**), **4-2** (**i**, **j**), **5-12** (**k**, **l**), and **6-1** (**m**, **n**). Experiments for **1-4** and **1-7** were performed in triplicate, with results given as mean ± SD, whereas **2-4**, **2-5**, **4-2**, **5-12**, and **6-1** were tested once and reported as exact values.

**Figure 3.**
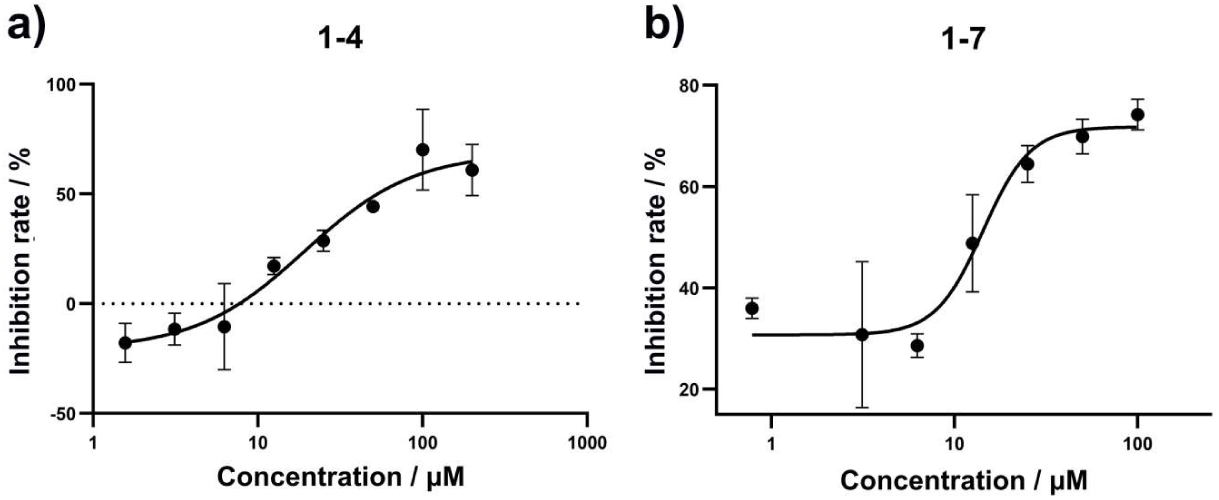
AlphaLISA-based inhibition assay. Inhibition curve of **1-4** (**a**) and **1-7** (**b**) against CHI3L1/galectin-3 interaction using the AlphaLISA inhibition assay. The inhibition rates were measured in triplicate with results given as the mean ± SD.

Although all four candidates were initially confirmed as CHI3L1 binders in the affinity measurements, considering the binding affinity and specificity, **2-4** and **2-5** were excluded from following steps, yielding a final hit rate of 0.28%. Following triplicate concentration-dependent assay, the *K*_d_ values of **1-4** and **1-7** were determined as 10.4 ± 1.0 μM and 7.40 ± 0.78 μM, respectively.

Interestingly, the predictions from the primary screening were highly consistent with the hits subsequently validated, highlighting the feasibility of applying the three sensorgram parameters for hit identification in HTS.

### AlphaLISA-based Binding Affinity Measurement

Galectin-3 is a β-galactoside-binding protein that interacts with CHI3L1 and plays an important role in the pathological processes of carcinogenesis, such as immune suppression, M2 macrophage polarization, and angiogenesis.^18^ In this study, an AlphaLISA assay was established to evaluate the inhibitory activities of **1-4** and **1-7** against the CHI3L1/galectin-3 interaction.

Compounds **1-4** and **1-7** (n = 3) were incubated with CHI3L1-His, galectin-3-GST, anti-His-donor beads, and anti-GST-acceptor beads in the dark, and the inhibition rate was measured based on a fluorescence method. Both compounds exhibited concentration-dependent inhibition of the CHI3L1/galectin-3 interaction, with IC_50_ values of 23.5 ± 2.0 μM and 15.4 ± 4.70 μM for **1-4** and **1-7**, respectively.

### Computational Study

Molecular docking of the hits **1-4** and **1-7** was carried out to investigate the binding mode of the hits. Both compounds **1-4** and **1-7** demonstrate favorable binding interactions with CHI3L1 (PDB ID: 8R4X)^21^ and share the binding pocket though they exhibit distinct binding modes and dynamic behaviors. **1-4** (**Figure 4a** - **4b**) exhibited a hydrogen bond with Asp207 and several hydrophobic interactions with Trp99, Met204, Met350 and Trp352 amino acid residues of the CHI3L1 binding site. Similarly, **1-7** (**Figure 4c** - **4d**) established one hydrogen bond with Trp99 and several hydrophobic interactions with Tyr27, Ala177, Met204, Phe261 and Arg263 amino acid residues of the binding site cavity. The similar modes of binding exhibited by both compounds help to explain the similar biological activities shown by both hits.

**Figure 4.**
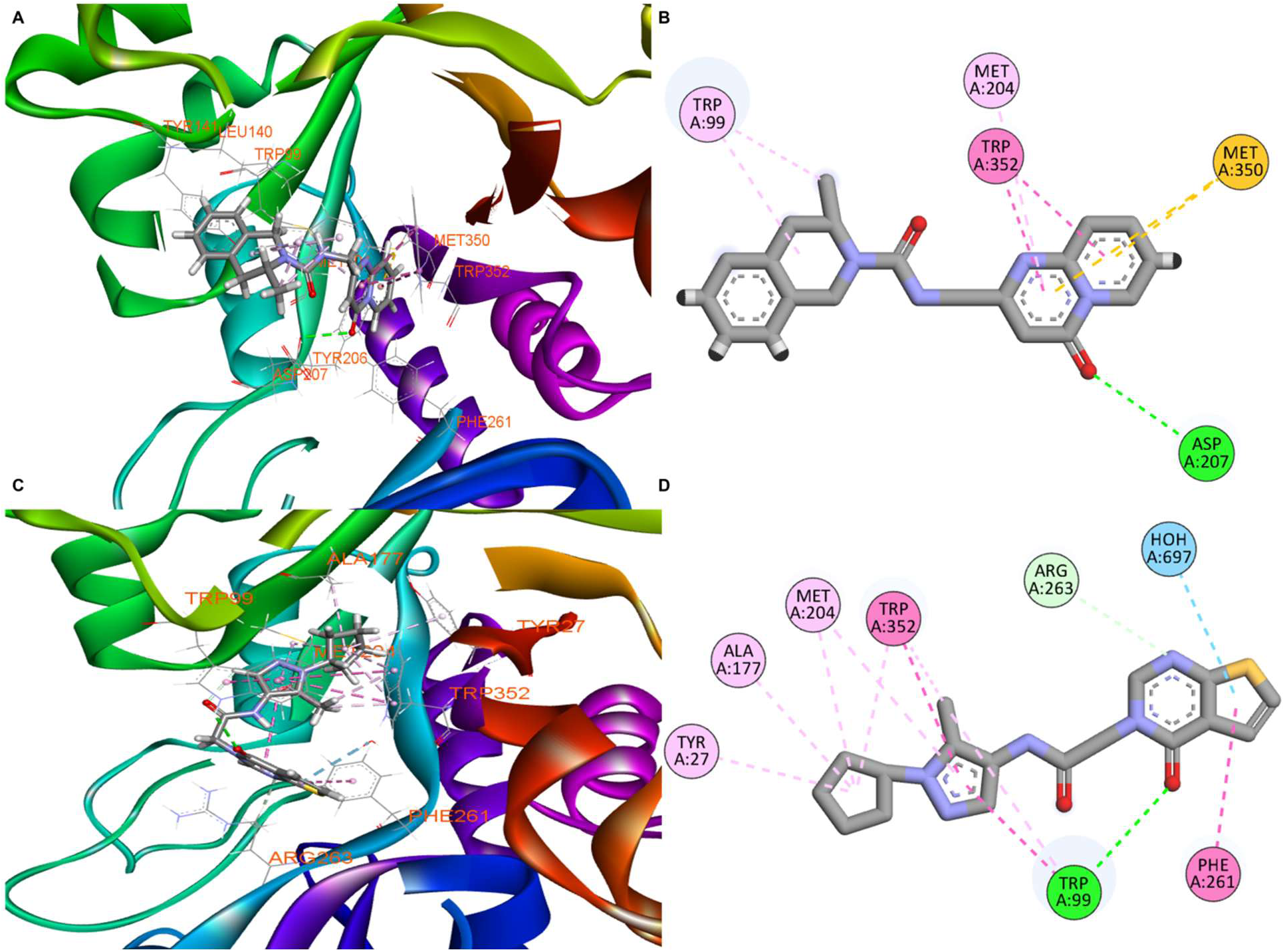
Molecular docking results of compounds 1-4 and 1-7 with CHI3L1. **(a)** 3D binding pose of compound **1-4** within the CHI3L1 binding pocket (PDB ID: 8R4X), showing key protein-ligand interactions. **(b)** 2D interaction diagram for compound **1-4**, illustrating hydrogen bonds (green dashed lines), π-π stacking interactions (pink dashed lines), and hydrophobic contacts with labeled residues. **(c)** 3D binding pose of compound **1-7** within the CHI3L1 binding pocket, displaying the alternative binding orientation and interaction network. **(d)** 2D interaction diagram for compound **1-7**, showing the distinct pattern of intermolecular interactions including hydrogen bonds, aromatic interactions, and hydrophobic contacts.

Next, three 100ns molecular dynamic simulations (MD) were carried out to validate the molecular docking results. The root mean square deviation (RMSD) analysis (**Figure 5a**) demonstrated that both ligand-bound systems maintained structural stability throughout the simulation period. The unbound CHI3L1 showed a RMSD of 1.15 Å. Meanwhile, both **1-4** and **1-7** exhibited an average RMSD of 1.1 and 1.12 Å, respectively. These results showed that CHI3L1 protein in complex with both hits induced stability. The RMSF analysis of the unbound CHI3L1 (**Figure 5b**) exhibited characteristic flexibility profiles with moderate fluctuations distributed across various regions, particularly showing elevated mobility in loop regions as expected. Upon binding of **1-4** (**Figure 5c**), CHI3L1 demonstrated altered flexibility patterns with some regions showing reduced mobility, likely due to stabilization effects from ligand binding. Notably, **1-7** binding (**Figure 5d**) resulted in more pronounced localized rigidity in the regions proximal to the binding site. This enhanced rigidity suggests that compound **1-7** may induce more significant conformational constraints, potentially contributing to its binding characteristics and biological activity profile compared to compound **1-4**.

**Figure 5.**
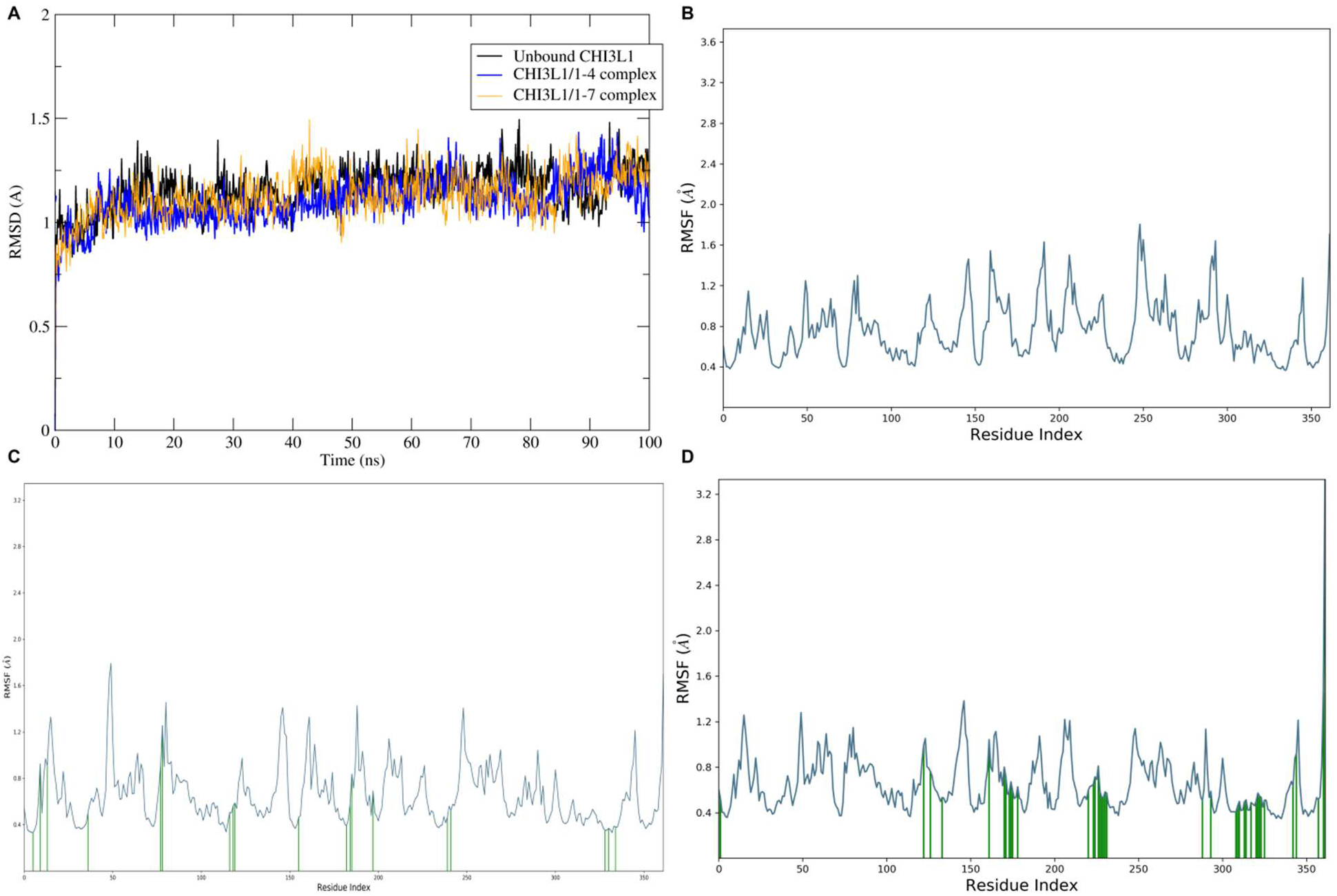
Molecular dynamics simulation analysis of CHI3L1 in different binding states. **(a)** Root mean square deviation (RMSD) over time showing structural stability of unbound CHI3L1 (black), CHI3L1/**1-4** complex (blue), and CHI3L1/**1-7** complex (orange) during 100 ns simulations. **(b)** Root mean square fluctuation (RMSF) per residue for unbound CHI3L1. **(c)** RMSF per residue for CHI3L1/**1-4** complex. **(d)** RMSF per residue for CHI3L1/**1-7** complex.

### Evaluation in GBM Spheroids

To investigate functional activity in a tumor-relevant setting, we evaluated compounds **1-4** and **1-7** using a multicellular 3D GBM spheroid model composed of GBM cells together with endothelial and macrophage populations. This spheroid system was selected because it more faithfully reproduces the complexity of the GBM microenvironment compared to conventional 2D cultures, allowing assessment of both cytotoxic and signaling outcomes. The previously described CHI3L1-STAT3 disruptor hygromycin B (HB) was included as a benchmark control.^25^

In this model, compound **1-7** demonstrated the most pronounced effects, producing a robust and concentration-dependent decrease in spheroid viability (**Figure 6a**). Significant reductions were observed already at 10 µM, with further loss of viability at 25 and 50 µM, indicating strong potency and consistent activity across doses. Compound **1-4** also reduced spheroid viability in a dose-responsive manner, although its overall effect was less pronounced than that of **1-7**. In contrast, HB showed only partial activity, achieving measurable reductions primarily at higher concentrations, thereby underscoring the superior efficacy of both novel compounds relative to the reference compound.

**Figure 6.**
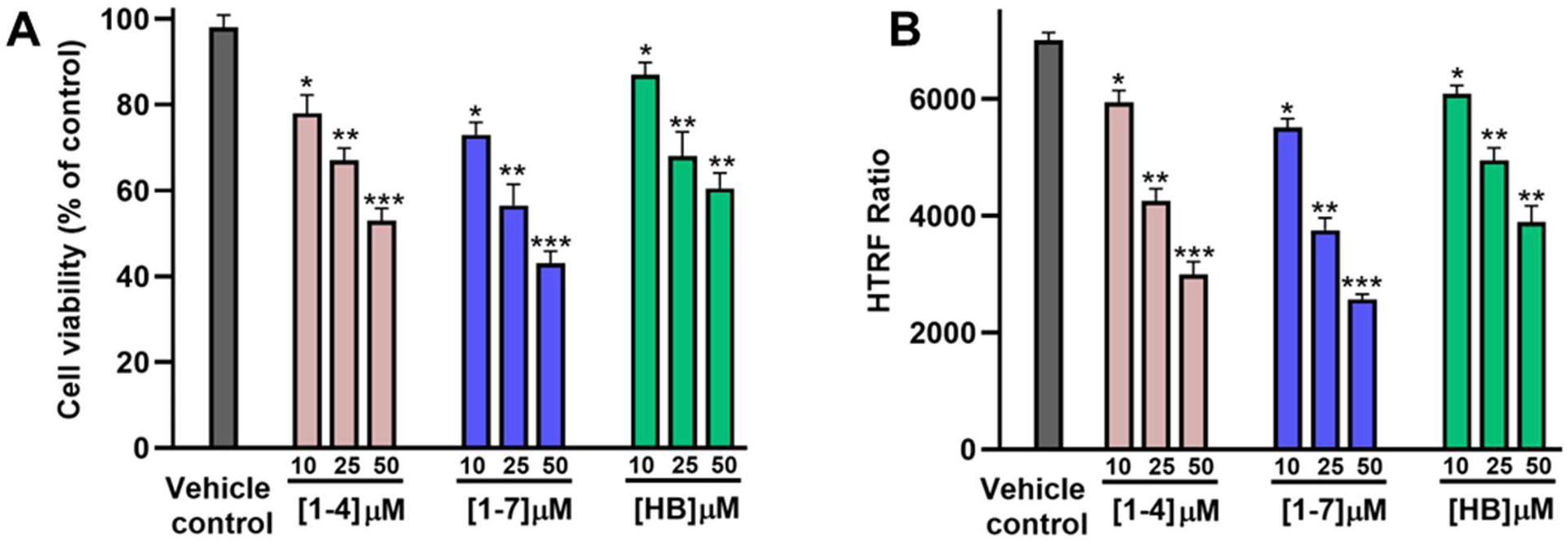
Evaluation of the therapeutic potential of compounds 1-4 and 1-7 in GBM spheroids. **(A)** Cell viability of GBM spheroids (as % of untreated control) upon incubation with increasing concentrations of the tested compounds (**1-4**, **1-7**, and HB) after 72 h incubation. **(B)** Reduction in phospho-STAT3 levels, as determined by HTRF phospho-STAT3 kit from Revvity (Cat# 62AT3PET), in GBM spheroids upon incubation with the tested compounds (**1-4**, **1-7**, and HB) after 72 h incubation. * *p* < 0.05, ** *p* < 0.01, and *** *p* < 0.001 relative to vehicle control. Data are representative of three independent experiments.

To further examine pathway engagement, we quantified levels of phosphorylated STAT3 (pSTAT3) using a homogeneous time-resolved fluorescence (HTRF) assay. As shown in **Figure 6b**, compound **1-7** produced marked inhibition of STAT3 phosphorylation across all concentrations tested, closely mirroring its effects on spheroid viability. Compound **1-4** exerted a moderate but significant suppression of pSTAT3, consistent with its intermediate binding affinity and biological activity. By comparison, HB again displayed weaker inhibitory activity, with less pronounced suppression of STAT3 signaling.

Together, these data establish compound **1-7** as the most potent CHI3L1 pathway modulator identified in this study, with superior ability to reduce GBM spheroid viability and inhibit STAT3 activation. Compound **1-4** also exhibited measurable efficacy, albeit to a lesser extent, while HB was comparatively weaker. These results validate the SPR-derived hits as functional inhibitors of CHI3L1 signaling in a physiologically relevant 3D model, providing a strong foundation for further optimization and in vivo studies.

## CONCLUSION

To identify novel molecular CHI3L1 binders, an SPR-based HTS platform was established and employed to screen a focus library of 704 molecules. In the primary screening at 100 μM, seven hits were identified with a relatively high occupancy rate (above 70%). Quantitative sensorgram analysis highlighted compounds **1-4** and **1-7** owing to their favorable binding kinetics, while other candidates were excluded due to slow dissociation. Following affinity measurement validated **1-4** and **1-7** as CHI3L1 binders with *K*_d_ of 10.4 ± 1.0 μM and 7.40 ± 0.78 μM, respectively, yielding a final hit rate of 0.28%. Their functional activities were further confirmed by AlphaLISA assays, where both compounds inhibited the CHI3L1/galectin-3 interaction with IC₅₀ values of 23.5 ± 2.0 μM and 15.4 ± 4.7 μM for **1-4** and **1-7**, respectively. Following computational study revealed a similar binding site and interaction model with CHI3L1 shared by both compounds, whereas **1-7** induced significant conformational constraints compared to **1-4**, contributing to the more potent biological properties than the latter. Importantly, both compounds demonstrated functional activity in a multicellular 3D GBM spheroid model, where **1-7** emerged as the most potent CHI3L1 pathway inhibitor, reducing spheroid viability and suppressing STAT3 activation more effectively than the reference compound HB. Collectively, these results not only establish the feasibility of SPR-based screening for non-enzymatic targets like CHI3L1 but also highlight compound **1-7** as a promising lead scaffold for further optimization and potential therapeutic development.

## EXPERIMENTAL

### SPR-based screening

The HTS was performed on the Biacore™ 8K SPR system (Cytiva, Marlborough, MA, USA), and 704 molecules from the Discovery Diversity Set (DDS-10, Enamine, Kyiv, Ukraine) were screened.

hCHI3L1-His Protein (8.0 μg/mL in acetate buffer, pH 5.0) was immobilized on a Series S Sensor Chip CM5 (29104988, Cytiva, Marlborough, MA, USA) using a commercial amine coupling kit (BR100050, Cytiva, Marlborough, MA, USA). Immobilization was performed at a flow rate of 10 μL/min for 420 s, reaching an immobilization level around 3000 RU, followed by blocking with ethanolamine. The flow cell with solely ethanolamine-block on the same channel served as the reference. Immobilization buffer: PBS-P+ (28995084, Cytiva, Marlborough, MA, USA).

2 μL of tested compounds were transferred into an intermediate 384-well plate using the MINI 96 channel portable electronic pipette (Integra Bioscience, Zizers, Switzerland), and 3 μL DMSO was added subsequently. After fully mixing, 3 μL of the above solution was added into 117 μL PBS-P+, and the final concentration of compound was 100 μM. The PBS-P+ solution of tested candidates were injected over the sensor chip in a single-injection model. The injection was performed at a flow rate of 30 μL/min for 120 s, followed by a regeneration step using regeneration buffer (10 mM sodium acetate, 250 mM NaCl, pH 4.0) with a flow rate of 30 μL/min for 30 s. Assay buffer: Assay buffer: PBS-P+ supplemented with 2.5% DMSO.

Data were analyzed using Biacore™ Insight Evaluation Software (Cytiva, Marlborough, MA, USA). The screening was performed in a single time.

### SPR-based Affinity Measurement

Gradient concentrations of compound were prepared in the assay buffer, and injected over the sensor chip in a single-cycle kinetics model with a flow rate at 30 μL/min for 120 s per injection. After each injection, a 30 s-regeneration was performed at a flow rate of 30 μL/min using the regeneration buffer.

Data were analyzed using Biacore™ Insight Evaluation Software (Cytiva, Marlborough, MA, USA). The assays were performed in a single time experiment and the *K*_d_ values were given as the exact numberm or in individual triplicate and the *K*_d_ values were given as mean ± SD.

### AlphaLISA-based Inhibitory Assay

Gradient concentrations of tested compound were prepared in the assay buffer (AL018C, Revvity, Inc., Waltham, M.A., U.S.) supplemented with 2.5% DMSO, and were incubated with hCHI3L1-His (CH1-H5228, Acro Biosystems, Newark, DE, USA) and Galectin-3 (10289-H09E, Sino Biological, Beijing, China), and the final concentrations of two proteins were both 60 nM. After incubating at room temperature (r.t.) for 3 h in dark, AlphaScreen anti-6His donor beads (15 μg/mL, AS116D, Revvity, Inc., Waltham, M.A., U.S) and AlphaLISA anti-GST acceptor beads (15 μg/mL, AL110C, Revvity, Inc., Waltham, M.A., U.S) were added, and an extra hour-incubation was performed at r.t. in dark. The signal detection was performed using a Tecan Spark. plate reader (Tecan Group Ltd., Männedorf, Switzerland) using the AlphaLISA model (wavelength: 623 nm; bandwidth: 25 nm). Assay buffer (2.5% DMSO) with CHI3L1-His and galectin-3-GST was performed as the positive control, while buffer with CHI3L1-His solely as the negative control. The inhibition rate was calculated as: inhibition rate = (fluorescence_sample_ - fluorescence_positive_) / (fluorescence_negative_ - fluorescence_positive_) × 100%. The assay was performed in technical triplicate and results were given as mean ± SD.

### Computational Study

Molecular docking studies were performed using Maestro Schrödinger (V.2023.4) suite to investigate the binding interactions of compounds **1-4** and **1-7** with CHI3L1 (PDB ID: 8R4X)^21^. The protein structure was prepared using the Protein Preparation Wizard, which included optimization of hydrogen bond networks, removal of water molecules, and energy minimization using the OPLS4 force field. Ligand structures were prepared using the LigPrep module with ionization states generated at pH 7.4 ± 2.0 using Epik. The binding site was defined based on the co-crystallized native ligand position, and receptor grid generation was performed using default parameters. Ligand docking was carried out using Glide with standard precision (SP) scoring function, and the best poses were selected based on docking score and visual inspection of binding interactions.

Following molecular docking, molecular dynamics (MD) simulations were conducted using DESMOND to validate the stability and dynamic behavior of the protein-ligand complexes. Three independent 100 ns MD simulations were performed for unbound CHI3L1, CHI3L1/**1-4** complex, and CHI3L1/**1-7** complex using appropriate MD simulation software. The systems were solvated in explicit water models and neutralized with counter ions. Energy minimization was performed followed by equilibration in NVT and NPT ensembles before production runs. Trajectory analysis was conducted to calculate root mean square deviation (RMSD) and root mean square fluctuation (RMSF) values. RMSD plots were generated using XMGRACE to visualize the structural stability of the systems over the simulation period. Three-dimensional visualization and analysis of protein-ligand interactions were performed using Discovery Studio Visualizer.

### GBM Spheroids Assay

The GBM spheroids were prepared as previously described.^26^ U-87 MG GBM cells (ATCC, Cat# HTB-14) were cultured in DMEM (ATCC, Cat#30-2002) containing 4.5 g/L glucose and 2 mM L-glutamine, supplemented with streptomycin and penicillin. HMEC-1 (ATCC) were maintained in MCDB131 medium with 10 mM L-glutamine, 10 ng/ml FGF, and 1 µg/ml hydrocortisone. Briefly, U-87 MG and HMEC-1 cells were co-seeded with macrophages on low-adhesion 96-well plates at 2 × 10³ cells per well in 100 µl of the tested compounds at varying concentrations (10, 25, and 50 µM) or control media. After 72 hours, cell viability was assessed using the CCK-8 assay (MedChemExpress, Cat# HY-K0301) according to the manufacturer’s recommended protocol. Absorbance at 450 nm was measured using a microplate reader.

Assessment of phospho-STAT3 levels was determined by HTRF phospho-STAT3 kit from Revvity (Cat# 62AT3PET) using the manufacturer’s recommended protocol. All experiments were conducted in triplicate.

## Acknowledgments

This work was supported by the National Institute of Neurological Disorders and Stroke under grant number R01NS136524 (PI: Gabr).

## Data availability

The authors confirm that all data are available as ESI.† Furthermore, additional data and original files are available from the authors upon request.

## Conflicts of interest

There are no conflicts to declare

## Notes

### Competing Interest Statement

The authors have declared no competing interest.

## REFERENCES

(1) J. Yu, I. Yeo, S. Han, J. Yun, B. Kim, Y. Yong, Y. Lim, T. Kim, D. Son, J. Hong, Significance of chitinase-3-like protein 1 in the pathogenesis of inflammatory diseases and cancer. Exp Mol Med 2024, 56 (1), 1–18.

(2) C. He, C. Lee, C. Dela Cruz, C. Lee, Y. Zhou, F. Ahangari, B. Ma, E. Herzog, S. Rosenberg, Y. Li, et al. Chitinase 3-like 1 regulates cellular and tissue responses via IL-13 receptor alpha2. Cell Rep 2013, 4 (4), 830–841.

(3) B. Lananna, C. McKee, M. King, J. Del-Aguila, J. Dimitry, F. Farias, C. Nadarajah, D. Xiong, C. Guo, A. Cammack, et al. Chi3l1/YKL-40 is controlled by the astrocyte circadian clock and regulates neuroinflammation and Alzheimer&#x2019;s disease pathogenesis. Science Translational Medicine 2020, 12 (574), eaax3519.

(4) T. Zhao, Z. Su, Y. Li, X. Zhang, Q. You, Chitinase-3 like-protein-1 function and its role in diseases. Signal Transduct Target Ther 2020, 5 (1), 201.

(5) I. Schmidt, I. Hall, S. Kale, S. Lee, C. He, Y. Lee, G. Chupp, G. Moeckel, C. Lee, J. Elias, et al. Chitinase-like protein Brp-39/YKL-40 modulates the renal response to ischemic injury and predicts delayed allograft function. J Am Soc Nephrol 2013, 24 (2), 309–319.

(6) C. Dela Cruz, W. Liu, C. He, A. Jacoby, A. Gornitzky, B. Ma, R. Flavell, C. Lee, J. Elias, Chitinase 3-like-1 promotes Streptococcus pneumoniae killing and augments host tolerance to lung antibacterial responses. Cell Host Microbe 2012, 12 (1), 34–46.

(7) N. Xu, Q. Bo, R. Shao, J. Liang, Y. Zhai, S. Yang, F. Wang, X. Sun, Chitinase-3-Like-1 Promotes M2 Macrophage Differentiation and Induces Choroidal Neovascularization in Neovascular Age-Related Macular Degeneration. Invest Ophthalmol Vis Sci 2019, 60 (14), 4596–4605.

(8) J. Bao, Y. Ouyang, L. Qiao, J. He, F. Liu, Y. Wang, L. Miao, A. Fu, Z. Lou, Q. Zang, et al. Serum CHI3L1 as a Biomarker for Non-invasive Diagnosis of Liver Fibrosis. Discov Med 2022, 33 (168), 41–49.

(9) H. Shantha Kumara, D. Gaita, H. Miyagaki, X. Yan, S. Hearth, L. Njoh, V. Cekic, R. Whelan, Plasma chitinase 3-like 1 is persistently elevated during first month after minimally invasive colorectal cancer resection. World J Gastrointest Oncol 2016, 8 (8), 607–614.

(10) A. Rusak, I. Buzalewicz, M. Mrozowska, B. Wiatrak, K. Haczkiewicz-Lesniak, M. Olbromski, A. Kmiecik, E. Krzyzak, A. Pietrowska, J. Moskal, et al. Multimodal study of CHI3L1 inhibition and its effect on angiogenesis, migration, immune response and refractive index of cellular structures in glioblastoma. Biomed Pharmacother 2023, 161, 114520.

(11) L. Hai, D. Hoffmann, R. Wagener, D. Azorin, D. Hausmann, R. Xie, M. Huppertz, J. Hiblot, P. Sievers, S. Heuer, et al. A clinically applicable connectivity signature for glioblastoma includes the tumor network driver CHI3L1. Nat Commun 2024, 15 (1), 968.

(12) S. Wang, S. Chen, M. Jin, M. Hu, W. Huang, Z. Jiang, J. Yang, Y. Zhang, H. Wu, Y. Hu, et al. Diagnostic and prognostic value of serum Chitinase 3-like protein 1 in hepatocellular carcinoma. J Clin Lab Anal 2022, 36 (2), e24234.

(13) M. Eldaly, F. Metwally, W. Shousha, A. El-Saiid, S. Ramadan, Clinical Potentials of miR-576-3p, miR-613, NDRG2 and YKL40 in Colorectal Cancer Patients. Asian Pac J Cancer Prev 2020, 21 (6), 1689–1695.

(14) J. Yu, I. Yeo, D. Son, J. Yun, S. Han, J. Hong, Anti-Chi3L1 antibody suppresses lung tumor growth and metastasis through inhibition of M2 polarization. Mol Oncol 2022, 16 (11), 2214–2234.

(15) J. Johansen, L. Drivsholm, P. Price, I. Christensen, High serum YKL-40 level in patients with small cell lung cancer is related to early death. Lung Cancer 2004, 46 (3), 333–340.

(16) D. Kim, H. Park, S. Lim, J. Koo, H. Lee, J. Choi, J. Oh, S. Ha, M. Kang, C. Lee, et al. Regulation of chitinase-3-like-1 in T cell elicits Th1 and cytotoxic responses to inhibit lung metastasis. Nat Commun 2018, 9 (1), 503.

(17) C. Lee, C. He, A. Nour, Y. Zhou, B. Ma, J. Park, K. Kim, C. Dela Cruz, L. Sharma, M. Nasr, et al. IL-13Ralpha2 uses TMEM219 in chitinase 3-like-1-induced signalling and effector responses. Nat Commun 2016, 7, 12752.

(18) P. Yang, M. Yu, Y. Hou, C. Chang, S. Lin, I. Kuo, P. Su, H. Cheng, W. Su, Y. Shan, et al. Targeting protumor factor chitinase-3-like-1 secreted by Rab37 vesicles for cancer immunotherapy. Theranostics 2022, 12 (1), 340–361.

(19) C. Lee, D. Hartl, G. Lee, B. Koller, H. Matsuura, C. Da Silva, M. Sohn, L. Cohn, R. Homer, A. Kozhich, et al. Role of breast regression protein 39 (BRP-39)/chitinase 3-like-1 in Th2 and IL-13-induced tissue responses and apoptosis. J Exp Med 2009, 206 (5), 1149–1166.

(20) M. Kawada, H. Seno, K. Kanda, Y. Nakanishi, R. Akitake, H. Komekado, K. Kawada, Y. Sakai, E. Mizoguchi, T. Chiba, Chitinase 3-like 1 promotes macrophage recruitment and angiogenesis in colorectal cancer. Oncogene 2012, 31 (26), 3111–3123.

(21) W. Czestkowski, L. Krzeminski, M. Piotrowicz, M. Mazur, E. Pluta, G. Andryianau, R. Koralewski, K. Matyszewski, S. Olejniczak, M. Kowalski, et al. Structure-Based Discovery of High-Affinity Small Molecule Ligands and Development of Tool Probes to Study the Role of Chitinase-3-Like Protein 1. J Med Chem 2024, 67 (5), 3959–3985.

(22) Y. Lee, J. Yu, K. Kim, D. Lee, D. Son, H. Lee, J. Jung, N. Kim, Y. Ham, J. Yun, et al. A small molecule targeting CHI3L1 inhibits lung metastasis by blocking IL-13Ralpha2-mediated JNK-AP-1 signals. Mol Oncol 2022, 16 (2), 508–526.

(23) H. Ham, Y. Lee, J. Yun, D. Son, H. Lee, S. Han, J. Hong, K284-6111 alleviates memory impairment and neuroinflammation in Tg2576 mice by inhibition of Chitinase-3-like 1 regulating ERK-dependent PTX3 pathway. J Neuroinflammation 2020, 17 (1), 350.

(24) D. Hong, J. Yu, J. Lee, D. Son, H. Lee, Y. Kim, J. Chang, D. Lee, W. Lee, J. Yun, et al. A Natural CHI3L1-Targeting Compound, Ebractenoid F, Inhibits Lung Cancer Cell Growth and Migration and Induces Apoptosis by Blocking CHI3L1/AKT Signals. Molecules 2022, 28 (1).

(25) W. Yu, S. Gui, L. Peng, H. Luo, J. Xie, J. Xiao, Y. Yilamu, Y. Sun, S. Cai, Z. Cheng, Z. Tao, STAT3-controlled CHI3L1/SPP1 positive feedback loop demonstrates the spatial heterogeneity and immune characteristics of glioblastoma. Dev Cell 2025, 60(12), 1751.

(26) H. Nada, L. Zhang, B. Kaur, M. Gabr, CHI3L1-Targeted Small Molecules as Glioblastoma Therapies: Virtual Screening-Based Discovery, Biophysical Validation, Pharmacokinetic Profiling, and Evaluation in Glioblastoma Spheroids. Eur J Med Chem 2025, 297, 117960.

