## Supporting Information for "Discovery of Small Molecule CHI3L1 Inhibitors by SPR-Based High-Throughput Screening"

| **Contents** |  |
| --- | --- |
| Primary SPR screening results | S2 |

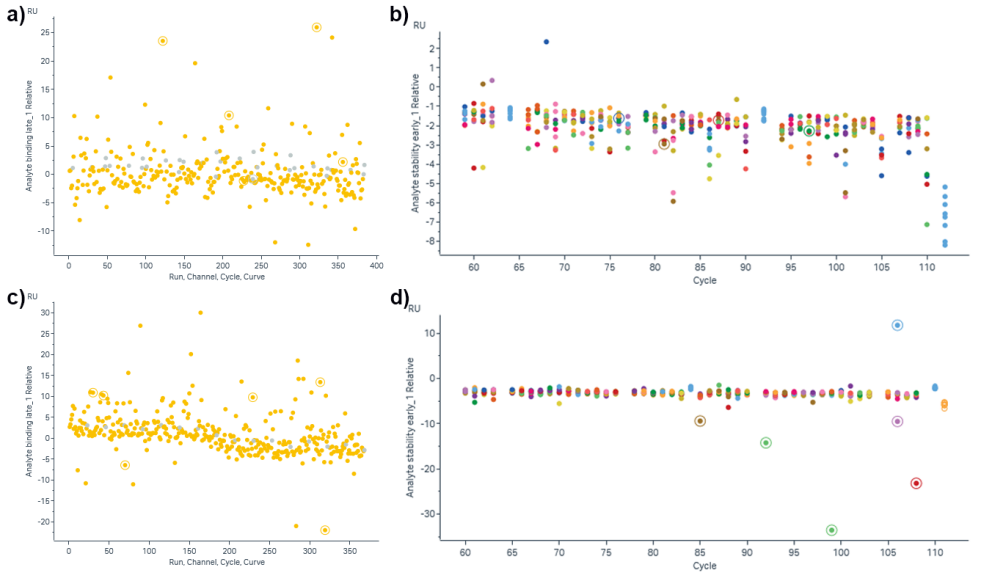

**Figure S1. Primary SPR screening results.** a and c) Relative response of candidates at 100 μM. b and d) Response signal of candidates in the reference cell.
